# New variants in the Chloroplast Genome Sequence of two Colombian individuals of the Cedar timber species (*Cedrela odorata* L.), using long read Oxford Nanopore Technology

**DOI:** 10.1101/2023.07.04.546705

**Authors:** Jaime Simbaqueba, Gina A. Garzón-Martínez, Nicolas Castano

**Affiliations:** Instituo Amazónico de Investigaciones Científicas SINCHI, Bogotá, Colombia; Centro de Investigación Tibaitatá, Corporación Colombiana de Investigación Agropecuaria (Agrosavia), Mosquera, Cundinamarca, Colombia; Nourish Ingredients, Canberra ACT, Australia

## Abstract

The plant species *Cedrela odorata* has been largely exploited in the timber industry due to the high demand of its wood. Therefore, *C. odorata*, has been considered as a vulnerable species since 1994, with the Convention on International Trade in Endangered Species of Wild Fauna and Flora (CITES). *C. odorata* is one of the key timber species included in the management and conservation plans in Colombia. These plans include the study of local populations by developing genetic resources for the species. In this study, two novel chloroplast (cp) genomes have been generated using the MinION long read sequencing technology. The new cp assemblies were compared with other cp genomes for the species, supporting previous hypothesis of intraspecific diversity associated to their place of origin. Furthermore, the cp genomes were screened for sequence variants (SV) and a total of 16 SNPs were identified, presumably unique to populations of the amazon region in Colombia. Here, the first cp genome sequences of *C. odorata* from Colombian individuals represent novel and valuable genomic resources for the species suitable for DNA fingerprinting and DNA barcoding applications.

## Introduction

*Cedrela odorata* L. is a widely distributed species from the *Meliaceae* family, that produces a valuable and multi-purpose timber. However, due to its continuous over-exploitation, it is listed as a vulnerable species on the International Union for Conservation of Nature (IUCN) Red List of Threatened Species in parts of the Americas (Mark & Rivers, 2017), and as a vulnerable species by the Convention on International Trade in Endangered Species of Wild Fauna and Flora (CITES) under Appendix II (Cites, 2021). Nowadays, illegal timber trade of *Cedrela* and other tree species is a widespread problem, responsible of deforestation, forest degradation, associated with biodiversity and economic impacts. And, despite the efforts of several countries to combat illegal timber trade, further actions in terms of policy, assessment of causes of illegal logging and novel strategies to control harvest and trade timber species need to be addressed (Bösch, 2021).

In Colombia, *C. odorata* can be found in different environmental gradients, from tropical dry forest to the pre montane and tropical moist forests (Cárdenas et al., 2015). However, natural populations in the country have been reduced in around 80% due to overexploitation, habitat fragmentation and the expansion of the agricultural border (Franco et al., 2019), to the point that cedar exploitation is not allowed from 2015 to 2025 in some areas (*Resolution 0110, 12 February*, 2015). Currently, this species is listed as vulnerable in Colombia by the IUCN and the Ministry of Environment and Sustainable Development under Resolution 1912, 2017. However, illegal trading with false documents of the species and geographical origin are still used for commercial purposes (Molinares et al., 2019).

To fight against illegal logging, new cost-effective and efficient tools need to be implemented. Genetic fingerprint is an ideal method to determine species identity and control the geographic origin of timber (Degen & Fladung, 2007). However, factors such as high-quality DNA, an affordable sequencing technology for the generation of suitable molecular markers, and the build of reliable reference data across species, determine the successful use of DNA barcoding for timber logging. Different molecular markers and strategies have been performed in timber species including *C. odorata*, to understand the population structure, genetic diversity, phylogenetic structure, harvesting origin, among others (Cavers et al., 2003, 2013; Degen & Fladung, 2007; J. L. Hu et al., 2022; Paredes-Villanueva et al., 2019).

Comparative genomics based on chloroplast DNA (cpDNA) sequences have become an ideal approach to be used as fingerprint (phylogenomics) and biogeographic analyses (B. Li et al., 2016; Shaw et al., 2014; Wei et al., 2017). The chloroplast genome (cp) is highly conserved in plants in terms of genes composition and arrangement, uniparentally inherited, and its abundance compared to the nuclear DNA allow the rapid distinction (i.e., assembly and annotation) of sequences, avoiding the need to purify the organelle (Schroeder et al., 2016). Cp data could increase the resolution and contribute to the development of new molecular markers to be used as specific barcodes for more than two loci (phylogenetics) (Hollingsworth et al., 2009; Mader et al., 2018).

Advances in sequencing technology allow to produce accurate and affordable data to study a wide range of species. Currently, third generation sequencing technology platforms can generate sequences > 10 kb (Giordano et al., 2017). Thus, compared to previous short read technologies, one or few reads could be sufficient to cover the entire cp genome offering a faster <u>turnaround time</u> with a great potential for population studies through single nucleotide polymorphism (SNP) genotyping (T. Hu et al., 2021; W. Wang et al., 2018). Long read Oxford Nanopore Technology (ONT) have been use in a diverse range of applications, including genome assembly (Michael et al., 2018), building new reference genomes (Jain et al., 2016; Scott et al., 2020), rapid diagnoses of plant viruses (Boykin et al., 2019), among others. The MinION, which uses the ONT technology, is a portable, pocket-sized device, ideal for species with few or null genomic information (Y. Wang et al., 2021), as most plant species of the Amazon rainforest, including the cedar timber *C. odorata*.

In this study, two individuals of *C. odorata* were selected to sequence their cp genome as a proof of concept, to establish a protocol for high molecular DNA extraction and perform a sequencing workflow using a cheap and fast portable sequencer MinION. As a result, the first genomic resources of *C. odorata* were obtained from two botanical samples of the Amazon region in Colombia. Subsequent phylogenomic analyses of the cp genomes corroborate the cryptic diversity demonstrated for the species and identified unique sequence variants (SV), that could be used in forensic genomics strategies aiming to combat the illegal timber trade in Colombia.

## Materials and Methods

### Plant material and High Molecular Weight (HMW) DNA extraction

Two *C. odorata* specimens were collected in the Colombian departments of Caquetá and Putumayo. Plant material is available at the Colombian Amazon Herbarium (COAH) with ID codes: OGP2096 and OGP2143. HMW DNA was extracted from fresh leaves, using the protocol reported for *Eucalyptus* species by Schalamun and Schwessinger (2017), with minor modifications. An additional clean-up steps with phenol:chlorophorm:Isoamyl alcohol (25:24:1) and isopropyl alcohol was added to ensure DNA quality. The DNA concentration was quantified by fluorometry Quibit^™^DNA kit and the quality was estimated with NanoDrop^™^ by comparing to the 260/280 and 260/230 nm ratios.

### MinION sequencing

Libraries were prepared according to the ligation library protocol LibPrep for SQK-LSK109 cells with no DNA fragmentation (Reiling et al., 2020). Long-read sequencing was carried out on a MinION Mk1C sequencer using the MinKNOW^™^ software release v20.10 (Jain et al., 2018). Data was base called using Albacore release v2.0.2 (https://github.com/Albacore/albacore). Quality control (QC) steps were performed according to Wang et al., (2018). Adaptors from long-reads were removed using Porechop v0.2.1 (https://github.com/rrwick/Porechop.) and bases with quality < 9 were trimmed using Nanofilt v1.2.0 (De Coster et al., 2018). Reads shorter than 5 kb were discarded.

### Chloroplast read extraction and assembly

The bioinformatics pipeline for chloroplast long-read extraction and assembly for *Eucalyptus* species was used, with some modifications (W. Wang et al., 2018). The long-reads data obtained from the whole genome sequencing strategy, previously described, were filtered for non-chloroplast sources, such as the nuclear and mitochondrial genome and other contaminants. Reads were aligned with the *C. odorata* reference chloroplast genome (GenBank accession NC_037251.1) (Mader et al., 2018), using BLASR software v5.1 (Chaisson & Tesler, 2012). A duplicated and concatenated reference genome was used to ensure the alignment of long-reads span the point at which the genome was circularised, according to the strategy used by Wang et al., (2018) for circular genomes. The assembly of the filtered reads for the OGP2096 specimen was performed using Canu v2.1.1 (Koren et al., 2017) and Hinge v0.4.1 software’s (Kamath et al., 2017), with default parameters, and assuming > 40X coverage of the reads to the reference genome. Later, the best performing assembler software was used for the OGP2143 specimen.

### Post-assembly polishing

The assembled contigs were concatenated using MuMmer v3.23 (Kurtz et al., 2004), and polished using Racon (Vaser et al., 2017) and Nanopolish v0.8.1 (Loman et al., 2015) software’s. Both assemblies were manually inspected for duplicate sequences with the software Geneious v. 2020.0.5.

### Variant discovery and phylogenetic analysis

A diversity panel of *Cedrela* species was generated by retrieving the chloroplast genome assemblies of 15 individuals, including the reference genome NC037251.1, an additional draft assemblies reported by Finch et al., (2019) and the chloroplast genomes of *C. odorata* generated in this study, as follows: 11 *C. odorata*, one *C. fissilis, C. montana, C angustifolia* and *C. saltensis* (Table 1). These assemblies were chosen for this analysis due to their close collection sites in Colombia or nearby. Sequence reads from the COAH specimens (OGP2096 and OGP2143), and draft assemblies were aligned to the *C. odorata* reference genome using BWA-MEM v. 0.7.17 (H. Li & Durbin, 2009). These alignments were used as inputs for probabilistic variant calling using the Snippy software from the Galaxy bioinformatics toolkit version (Afgan et al., 2018) and the Genome Analysis Toolkit (GATK) v4.2.4.0 (Van der Auwera & O’Connor, 2020) including high-stringency variant filtering for coverage, mapping quality, and variant position. SNP filtering was performed with VCFtools v.0.1.17 (Danecek et al., 2011), to remove insertion/deletion variants, sites with greater than 85% of missing data and more than two alleles. Manual inspection of the genomic regions with SNPs were performed by comparing their positions with the reference genome NC_037251.1, using Geneious v.2020.0.5 software (https://www.geneious.com).

**Table 1.** Chloroplast genomes of *Cedrela* genus used for phylogenomic analyses.

| Sequence name | Species | Collection site | Reference |
| --- | --- | --- | --- |
| CEAN143 | *C. angustifolia* | Southern Bolivia | Finch et al., (2019) |
| CEFI211 | *C. fissilis* | Northern Bolivia |  |
| CEMO50 | *C. montana* | Western Colombia |  |
| CESA102 | *C. saltensis* | Southern Bolivia |  |
| COD10 | *C. odorata* | Nicaragua |  |
| COD52 |  | Costa Rica |  |
| COD162 |  | Nicaragua |  |
| COD185 |  | Costa Rica |  |
| COD202 |  | Venezuela |  |
| COD222 |  | Panamá |  |
| COD277 |  | Costa Rica |  |
| NOVO_NYBG |  | Mexico |  |
| NC037251.1 | *C. odorata* | Cuba | Mader et al., (2018) |
| OGP2096 | *C. odorata* | Colombia (Caquetá) | This study |
| OGP2143 |  | Colombia (Putumayo) |  |

Phylogenetic relationships were estimated with 1000 bootstrap Bayesian probabilities using the software BEAST (Bayesian Evolutionary Analysis Sampling Trees) v 2.6.1 (Bouckaert et al., 2019), with default settings. The resulting phylogenetic trees were visualised with the Interactive Tree of Life (iTOL) v4 web-based software (Letunic & Bork, 2019).

## Results

### *C. odorata* whole genome sequencing

A total of 2.6 and 0.99 Gb of raw long-read data were generated for individuals OGP2096 and OGP2143, respectively (Table 2). OGP2096 had a total of 594,819 reads, compared to OGP2143 with 110,484 reads. However, longer reads were observed for OGP2143, with a mean read length of 9.6 kb, compared to OGP2096, with a mean length of 4.3 kb (Table 2, Figure 1). The mean quality values for both sequenced individuals were 10 and 9.9, respectively.

**Table 2.** *C. odorata* ONT raw data.

| Sample | No. Reads | Mean read length (kb) | Mean Quality value | Total sequences produced (Gb) |
| --- | --- | --- | --- | --- |
| **OGP2096** | 594,819 | 4.37 | 10 | 2.6 |
| **OGP2143** | 110,484 | 9.67 | 9.9 | 1.0 |

**Figure 1.**
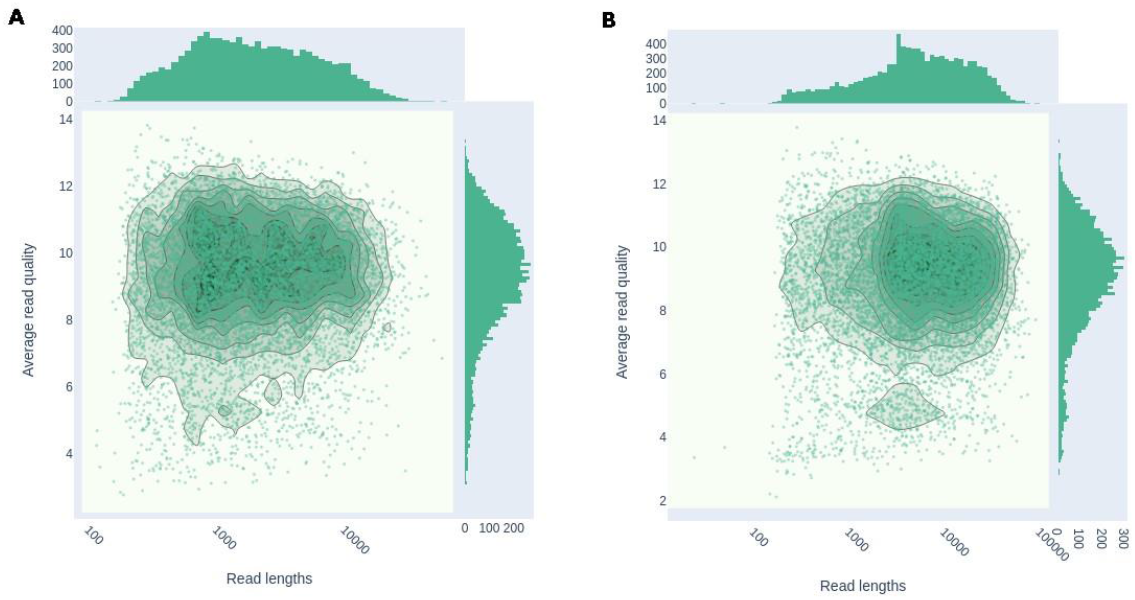
ONT quality sequencing analysis of raw data. Read lengths vs average read from **A**. OGP2096 and **B**. OGP2143.

### Chloroplast assembly

A total of 1,437 reads with a mean read length of ∼9,000 bp from the individual OGP2096 and 884 reads with a mean length of ∼11,000 bp from the individual OGP2143 were mapped to the *C. odorata* reference chloroplast genome (Table 3). Considering that the size of *C. odorata* cp genome is ∼ 159 kb, these subsets of reads represent a total coverage of ∼80X and ∼60X, respectively.

**Table 3.**
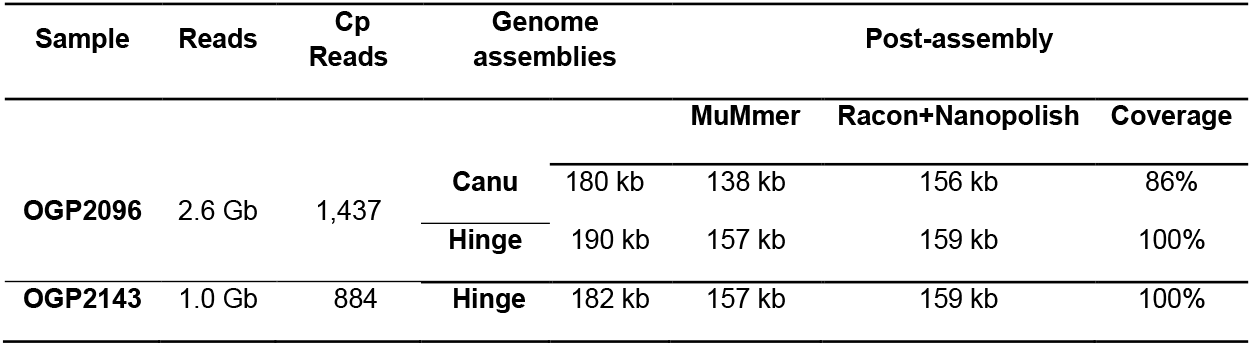
*C. odorata* cp genome assembly statistics

The subset of reads from OGP2096 were used to assemble this specimen cp genome using Canu. The resulting assembly was fragmented into three contigs of 102 kb (Contig 4), 55 kb (Contig 2) and 23 kb (Contig 3) in length (Supplementary Figure S1A). Contig 2 contains the large single copy region (LSC) and part of the inverted repeats (IR) of the chloroplast. Contig 4 contains part of the IRs and a small single copy (SSC), while Contig 3 contains part of the IR. These contigs were assembled in a unique contig of ∼138 kb by polishing duplicated regions (Table 3, Supplementary Figure S1B). This single contig is shorter in length compared to the *C. odorata* reference chloroplast genome (159 kb).

On the other hand, Hinge software showed an accurate performance for long-reads for the OGP2096 specimen, obtaining an assembly of 190 kb in length, fragmented into two contigs of 70 kb (Contig 1) and 120 kb (Contig 2) (Table 3, Supplementary Figure S1C), and a unique Contig of 157 kb after post assembly polishing and concatenation strategy (Table 3, Supplementary Figure S1D). Considering a better performance of the Hinge assembler, a total of 884 reads from the individual OGP2143 mapped to the *C. odorata* reference chloroplast genome, and they were used to assemble the OGP2143 cp genome. This assembly showed ∼182 kb in length and it was fragmented onto two contigs of 40 kb (Contig 1) and 142 kb (Conti 2) (Table 3, Supplementary Figure S1E). A unique contig of ∼157 kb after post assembly polishing analysis was obtained (Supplementary Figure S1F).

### Post assembly analysis

Racon and Nanopolish software were used to correct the above-mentioned genome assemblies. Furthermore, assemblies were manually inspected and compared with the *C. odorata* reference chloroplast genome, using the software Geneious v.2020.0.5. After removing duplicated regions, the size of the complete cp genome assembly was ∼159 kb in length for both specimens (Table 3). These assemblies were used in the prediction of the gene content and functional annotation of the cp genome (Figure 2). The cp genome assemblies of OGP2096 and OGP2143 specimens were uploaded into the GenBank under accession numbers OP750006 and OP750007, respectively.

**Figure 2.**
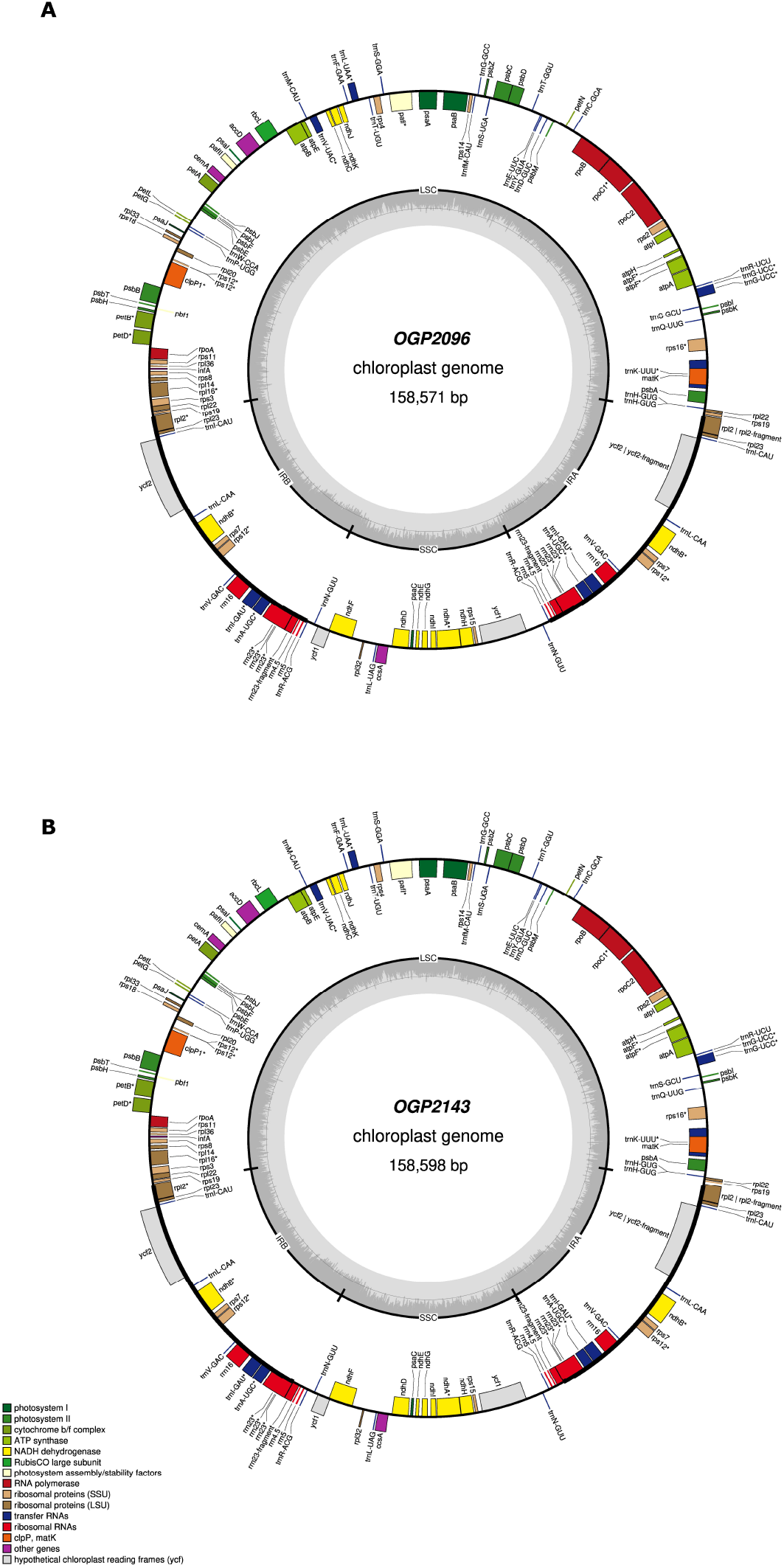
Graphic representation of the complete genetic map of the chloroplast of specimens **A**. OGP2096 and **B**. OGP2143 sequenced by the long-read Minion technology. GC content is represented by the inner grey circle. Genes inside the circle are transcribed clockwise, and those outside are transcribed counter clockwise. LSC, large single-copy; SSC, small single-copy; IR, inverted repeat.

### Phylogenetics and structural variation analysis

The phylogenetic analysis was performed by aligning the cp genome sequence of the *C. odorata* specimens OGP2096 and OGP2143 from Colombia, the cp reference sequence genome and sequences of the *Cedrela* genus (Table 1). The resulting phylogenetic trees support the monophyletic origin for all taxa of the *Cedrela* genus, as reported by Finch et al., (2019). The specimen OGP2096 grouped in a sister subclade of *Cedrela* species, closer to the out grouped *C. odorata* individual COD202 and other South American individuals (Subclade SA) (Figure 3A). The specimen OGP2143 is strongly grouped with the *C. monatana* individual CEMO50, forming a sister subclade close to mostly Central American (Subclade CA) individuals (Figure 3A).

**Figure 3.**
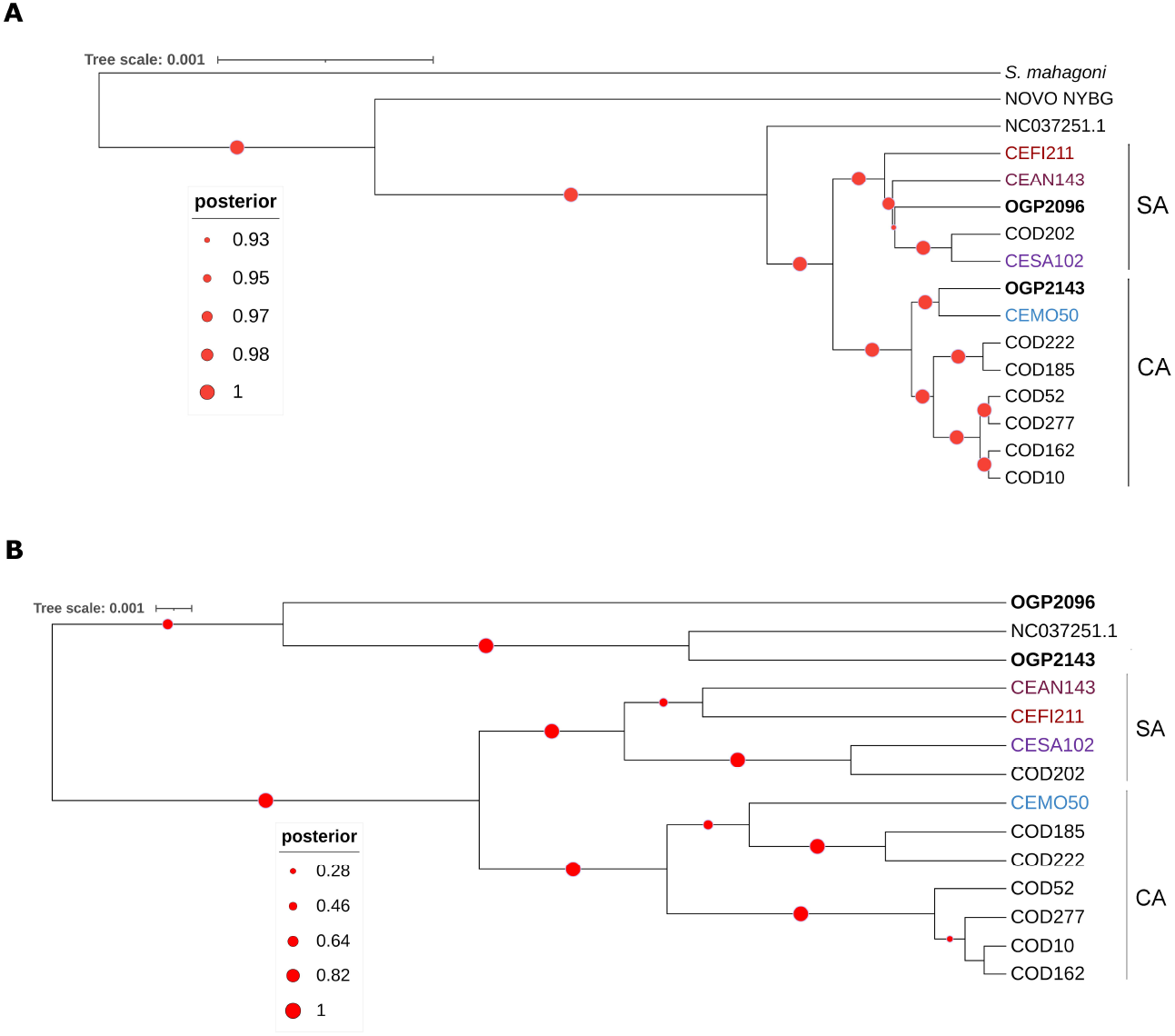
Phylogenetic analysis using the cp genome of individuals of the *Cedrela* genus. **A**. Tree generated with the alignment of complete cp genome sequences of all individuals. The analysis shows the grouping of two clades with significative bootstrap support, Central American Subclade (CA) and South American Subclade (SA), showing the intraspecific variation of the individuals. **B**. Resulted tree of the alignment of the sequence variants, identified with Snippy software of the Galaxy suit. The tree shows similar grouping according to the place or origin. The phylogenomic analyses were performed using BEAST phylogenetics software and the visualisation was carried out using the iTOL software. Black labelled taxa indicate *C. odorata* specimens collected in Colombia and deposited in the COAH. Coloured labelled taxa indicate other *Cedrela* species. The red circles indicate the bootstrap supporting as Bayesian posterior probability

Sequence variants were predicted to determine a group of SNPs that could be used to establish genotype studies in local populations of *C. odorata* from Colombia. A total of 393 deletions, 208 insertions and 96 multiple variants, were identified in the reference cp genome (NC_037251.1), compared to the 13 of the 14 cp genomes included in this study (Table 4). A total of 2,793 sequence variants (SNPs type), were also identified. From these set of SNPs, 187 were identified as unique of each individual compared to the reference. In addition, total of 131 SNPs segregates in the *Cedrela* species analysed (15 with *C. montana*, 20 to *C. saltensis*, 64 to *C. fissilis* and 32 with *C. angustifolia*, respectively). A few SNPs in the genomes of *C. odorata* were also observed (e.g., one for CEOD10, CEOD162 and two for OGP2143). The resulting phylogenetic analysis of *Cedrela* species based on the sequence variants, grouped the Colombian individuals sequenced in this study (OGP2096 and OGP2143), close to the reference genome (Figure 3B), perhaps reflecting the nearly complete genome assemblies generated with ONT long reads.

**Table 4.** Sequence variants identified in this study, using GATK and Galaxy pipelines.

| Individual | SNPs |  | INS | DEL | MNP | Total |
| --- | --- | --- | --- | --- | --- | --- |
|  | Unique | Total |  |  |  |  |
| CEAN143 | 32 | 249 | 23 | 39 | 5 | 348 |
| CEFI211 | 64 | 277 | 21 | 45 | 11 | 418 |
| CEMO50 | 15 | 221 | 13 | 28 | 6 | 283 |
| CESA102 | 20 | 270 | 20 | 44 | 9 | 363 |
| COD10 | 1 | 221 | 21 | 34 | 9 | 286 |
| COD52 | 2 | 239 | 18 | 36 | 11 | 306 |
| COD162 | 1 | 238 | 17 | 37 | 8 | 301 |
| COD185 | 9 | 240 | 19 | 31 | 7 | 306 |
| COD202 | 20 | 267 | 21 | 32 | 10 | 350 |
| COD222 | 6 | 228 | 19 | 28 | 7 | 288 |
| COD277 | 1 | 227 | 16 | 35 | 9 | 288 |
| OGP2096 | 14 | 70 | 0 | 2 | 3 | 89 |
| OGP2143 | 2 | 46 | 0 | 2 | 1 | 51 |
| **Total** | **187** | **2,793** | **208** | **393** | **96** | **3,677** |

INS= Insertions; DEL= Deletions; MNP = Multiplex SNPs

A total of 16 SV present in only OGP2096 and OGP2143 specimens were investigated by comparing the corresponding positions in the reference genome. In this analysis, six were identified as intergenic variants, six as intron variants and the remaining four correspond to missense variants for specific loci (Table 5).

**Table 5.** Potential *C. odorata*-specific SV for Colombian individuals identified from the chloroplast genome sequencing.

| NC_037251.1 |  | SNPs |  | Type | Locus |
| --- | --- | --- | --- | --- | --- |
| POS (nt) | REF | OGP2096 | OGP2143 |  |  |
| 6727 | G | A | G | variant | intergenic |
| 9279 | G | A | G | variant | intergenic |
| 16056 | C | T | C | variant | intergenic |
| 21511 | C | T | C | missense | rpoC2 |
| 40324 | T | A | T | missense | psaB |
| 44137 | G | A | G | variant | intergenic |
| 66416 | C | C | A | variant | intergenic |
| 75039 | G | A | G | variant | intergenic |
| 77654 | T | G | T | intron variant | psbH |
| 80024 | C | T | C | intron variant | petD |
| 84403 | G | A | G | intron variant* | Rpl16 |
| 87863 | G | G | A | intron variant | rpl2 |
| 114009 | C | T | C | missense | NdhF |
| 116740 | C | T | C | intron variant | Rpl32 |
| 124608 | G | A | G | intron variant | ndhA |
| 126930 | T | G | T | missense | rps15 |

\*Variant on splice donor

## Discussion

*Cedrela odorata* is an important plant species due to its use in the timber industry. However, because of the overexploitation of natural populations of *C. odorata* and related *Cedrela* species, the whole *Cedrela* genus has been declared as protected under CITES convention (Cites, 2021). Colombia is one of the countries with high biodiversity of *Cedrela* species and therefore, their wood is used in the timber industry, mainly illegal. Nevertheless, there is a lack of knowledge about natural populations of *C. odorata* and the locations where the timber is obtained. Thus, there is a need to establish suitable policies that limit the use of the natural resources of *C. odorata*.

Here, the chloroplast genome sequence of two individuals of *C. odorata* were generated using the emerging sequencing technology ONT: 1) As a proof of concept to obtain complete cp sequences as novel genomic resources for the species in Colombia, 2) These new resources were used in further phylogenetic analysis suggesting a possible intraspecific diversity (Figure 3), supporting the hypothesis of differences in local populations as reported by Finch et al, (2022), and 3) The new cp sequences were used to identify sequence variants that could be used in future tools developments to track timber resources of *C. odorata* collected illegally.

These are the first cp genomes of *C. odorata*, generated with the ONT long-read sequencing technology and the assembly pipeline for chloroplast genomes reported by Wang et al., (2018), for the wood forest species *Eucalyptus pauciflora*. Although the amount of raw long read data was low compared to other genome sequencing studies in plants (J. Wang et al., 2020; Wöhner et al., 2021) the number of raw reads (2.6 Gb and ∼1 Gb for the individuals OGP2096 and OGP2143, respectively), was sufficient to cover the cp genome up to 40X and generate complete cp genome assemblies for both individuals. These results also showed that assemblies generated by Hinge with further polishing steps were more accurate than Canu as demonstrated for *E. pauciflora* (W. Wang et al., 2018), thus suggesting the pipeline developed to assemble chloroplast genomes from long-read sequencing projects could be used in different plant species.

Plastid sequences of *C. odorata* can be used to support the geographic identification of derived timber material (Cavers et al., 2013; Finch et al., 2019). This phylogenetic analysis included comparison of both individuals OG2096 and OGP2143 with a subsample of fragmented cp genomes of *C. odorata* and *Cedrela* species, reported by Finch et al.,(2019), and a complete reference genome for the species (Figure 3A). The grouping of these *C. odorata* individuals, closely related to other *Cedrela* individuals from Colombia, corroborate the results of Finch et al., (2019) and Cavers et al., (2013), suggesting that cp genomes of *C. odorata* could be used to predict biogeography patterns on *C. odorata* individuals. Thus, these new chloroplast genomes could be used to identify broad scale local populations of *C. odorata* in Colombia. Moreover, recent evidence based on nuclear SNPs analysis of different species of *Cedrela*, suggest a cryptic diversity for *C. odorata*, differentiating Mesoamerican from South American individuals (Finch et al., 2022). Here, the phylogenetic analysis of the new cp genomes (Figure 3), could support the hypothesis of cryptic diversity for *C. odorata* reported by Finch et al., (2022), as OGP2143 are grouped with the Mesoamerican individuals while OGP2096 is more related to South American individuals. This hypothesis will be corroborated in future SNPs analysis of local populations of *C. odorata* from Colombia.

High throughput sequencing technologies allow the identification of SV that could be adapted as molecular markers for plant identification (Amarasinghe et al., 2020; Jung et al., 2019). Here, sequence variants between *C. odorata* individuals were identified, focusing on unique SV of OGP2096 and OGP2143 (Table 4). These SV are located on highly variable genomic regions of the *C. odorata* chloroplast, compared to other individuals of *Meliaceae* family (Mader et al., 2019). The intergenic (C/A) and rpl2 intron variants (G/A) in OGP2143 (Table 5), could be included in genotyping studies of *C. odorata* natural populations, in order to implement a genomic fingerprint for individuals extracted from the Amazon Region of Colombia.

## Supplementary material

**Figure S1**. MuMmer plot of the chloroplast assembly from **A**. OGP2096 cp genome organisation using Canu and **B**. its concatenated and polished assembly; **C**. OGP2096 cp genome assembly using Hinge and **D**. its concatenated and polished assembly; **E**. OGP2143 cp genome assembly using Hinge and **F**. its concatenated and polished assembly. Violet and red lines represent the length in kb of each assembled contig. Blue lines represent the genomic regions annotated in the *C. odorata* reference chloroplast genome (GenBank accession NC_037251.1)

## References

Afgan, E., Baker, D., Batut, B., Van Den Beek, M., Bouvier, D., Ech, M., Chilton, J., Clements, D., Coraor, N., Grüning, B. A., Guerler, A., Hillman-Jackson, J., Hiltemann, S., Jalili, V., Rasche, H., Soranzo, N., Goecks, J., Taylor, J., Nekrutenko, A., & Blankenberg, D. (2018). The Galaxy platform for accessible, reproducible and collaborative biomedical analyses: 2018 update. Nucleic Acids Research, 46(W1), W537–W544. https://doi.org/10.1093/NAR/GKY379

Amarasinghe, S. L., Su, S., Dong, X., Zappia, L., Ritchie, M. E., & Gouil, Q. (2020). Opportunities and challenges in long-read sequencing data analysis. In Genome Biology (Vol. 21, Issue 1). BioMed Central Ltd. https://doi.org/10.1186/s13059-020-1935-5

Bösch, M. (2021). Institutional quality, economic development and illegal logging: a quantitative cross-national analysis. European Journal of Forest Research, 140(5), 1049–1064. https://doi.org/10.1007/S10342-021-01382-Z/TABLES/3

Bouckaert, R., Vaughan, T. G., Barido-Sottani, J., Duchêne, S., Fourment, M., Gavryushkina, A., Heled, J., Jones, G., Kühnert, D., De Maio, N., Matschiner, M., Mendes, F. K., Müller, N. F., Ogilvie, H. A., Du Plessis, L., Popinga, A., Rambaut, A., Rasmussen, D., Siveroni, I., … Drummond, A. J. (2019). BEAST 2.5: An advanced software platform for Bayesian evolutionary analysis. PLOS Computational Biology, 15(4), e1006650. https://doi.org/10.1371/JOURNAL.PCBI.1006650

Boykin, L. M., Sseruwagi, P., Alicai, T., Ateka, E., Mohammed, I. U., Stanton, J.-A. L., Kayuki, C., Mark, D., Fute, T., Erasto, J., Bachwenkizi, H., Muga, B., Mumo, N., Mwangi, J., Abidrabo, P., Okao-Okuja, G., Omuut, G., Akol, J., Apio, H. B., … Ndunguru, J. (2019). Tree Lab: Portable Genomics for Early Detection of Plant Viruses and Pests in Sub-Saharan Africa. Genes, 10, 632. https://doi.org/10.3390/genes10090632

Cavers, S., Navarro, C., & Lowe, A. J. (2003). Chloroplast DNA phylogeography reveals colonization history of a Neotropical tree, Cedrela odorata L., in Mesoamerica. Molecular Ecology, 12(6), 1451–1460. https://doi.org/10.1046/J.1365-294X.2003.01810.X

Cavers, S., Telford, A., Arenal Cruz, F., Pérez Castañeda, A. J., Valencia, R., Navarro, C., Buonamici, A., Lowe, A. J., & Vendramin, G. G. (2013). Cryptic species and phylogeographical structure in the tree Cedrela odorata L. throughout the Neotropics. Journal of Biogeography, 40(4), 732–746. https://doi.org/10.1111/JBI.12086

Chaisson, M. J., & Tesler, G. (2012). Mapping single molecule sequencing reads using basic local alignment with successive refinement (BLASR): Application and theory. BMC Bioinformatics, 13(1), 1–18. https://doi.org/10.1186/1471-2105-13-238/FIGURES/11

Cites. (2021). CITES Appendices I, II, and III. Journal of Minimal Access Surgery, 78.

Resolution 0110, 12 February, (2015) (testimony of CORPOAMAZONIA).

Danecek, P., Auton, A., Abecasis, G., Albers, C. A., Banks, E., DePristo, M. A., Handsaker, R. E., Lunter, G., Marth, G. T., Sherry, S. T., McVean, G., Durbin, R., & Group, 1000 Genomes Project Analysis. (2011). The variant call format and VCFtools. Bioinformatics, 27(15), 2156–2158. https://doi.org/10.1093/BIOINFORMATICS/BTR330

De Coster, W., D’Hert, S., Schultz, D. T., Cruts, M., & Van Broeckhoven, C. (2018). NanoPack: visualizing and processing long-read sequencing data. Bioinformatics, 34(15), 2666–2669. https://doi.org/10.1093/BIOINFORMATICS/BTY149

Degen, B., & Fladung, M. (2007). Use of DNA-markers for tracing illegal logging. Proceedings of the International Workshop Fingerprinting Methods For, 321, 6–14.

Finch, K. N., Jones, F. A., & Cronn, R. C. (2019). Genomic resources for the Neotropical tree genus Cedrela (Meliaceae) and its relatives. BMC Genomics, 20(1). https://doi.org/10.1186/s12864-018-5382-6

Finch, K. N., Jones, F. A., & Cronn, R. C. (2022). Cryptic species diversity in a widespread neotropical tree genus: The case of Cedrela odorata. American Journal of Botany, 109(10), 1622–1640. https://doi.org/10.1002/ajb2.16064

Franco, N., Clavijo, C., Rojas, J., & Talero, C. (2019). Plan de Manejo y Conservación del Cedro (Cedrela odorata L.) para la jurisdicción de la Corporación Autónoma Regional de Cundinamarca CAR.

Giordano, F., Aigrain, L., Quail, M. A., Coupland, P., Bonfield, J. K., Davies, R. M., Tischler, G., Jackson, D. K., Keane, T. M., Li, J., Yue, J.-X., Liti, G., Durbin, R., & Ning, Z. (2017). De novo yeast genome assemblies from MinION, PacBio and MiSeq platforms. Scientific Reports, 3935. https://doi.org/10.1038/s41598-017-03996-z

Hollingsworth, P. M., Forrest, L. L., Spouge, J. L., Hajibabaei, M., Ratnasingham, S., van der Bank, M., Chase, M. W., Cowan, R. S., Erickson, D. L., Fazekas, A. J., Graham, S. W., James, K. E., Kim, K. J., John Kress, W., Schneider, H., van AlphenStahl, J., Barrett, S. C. H., van den Berg, C., Bogarin, D., … Little, D. P. (2009). A DNA barcode for land plants. Proceedings of the National Academy of Sciences of the United States of America, 106(31), 12794–12797. https://doi.org/10.1073/PNAS.0905845106/

Hu, J. L., Ci, X. Q., Liu, Z. F., Dormontt, E. E., Conran, J. G., Lowe, A. J., & Li, J. (2022). Assessing candidate DNA barcodes for Chinese and internationally traded timber species. Molecular Ecology Resources, 22(4), 1478–1492. https://doi.org/10.1111/1755-0998.13546

Hu, T., Chitnis, N., Monos, D., & Dinh, A. (2021). Next-generation sequencing technologies: An overview. Human Immunology, 82(11), 801–811. https://doi.org/10.1016/J.HUMIMM.2021.02.012

Jain, M., Koren, S., Miga, K. H., Quick, J., Rand, A. C., Sasani, T. A., Tyson, J. R., Beggs, A. D., Dilthey, A. T., Fiddes, I. T., Malla, S., Marriott, H., Nieto, T., O’Grady, J., Olsen, H. E., Pedersen, B. S., Rhie, A., Richardson, H., Quinlan, A. R., … Loose, M. (2018). Nanopore sequencing and assembly of a human genome with ultra-long reads. Nature Biotechnology, 36(4), 338–345. https://doi.org/10.1038/nbt.4060

Jain, M., Olsen, H. E., Paten, B., & Akeson, M. (2016). The Oxford Nanopore MinION: delivery of nanopore sequencing to the genomics community. Genome Biology, 17(1), 1–11. https://doi.org/10.1186/s13059-016-1103-0

Jung, H., Winefield, C., Bombarely, A., Prentis, P., & Waterhouse, P. (2019). Tools and Strategies for Long-Read Sequencing and De Novo Assembly of Plant Genomes. Trends in Plant Science, 24(8), 700–724. https://doi.org/10.1016/J.TPLANTS.2019.05.003

Kamath, G. M., Shomorony, I., Xia, F., Courtade, T. A., & Tse, D. N. (2017). HINGE: Long-read assembly achieves optimal repeat resolution. Genome Research, 27(5), 747–756. https://doi.org/10.1101/GR.216465.116/-/DC1

Koren, S., Walenz, B. P., Berlin, K., Miller, J. R., Bergman, N. H., & Phillippy, A. M. (2017). Canu: scalable and accurate long-read assembly via adaptive k-mer weighting and repeat separation. Genome Research, 27(5), 722–736. https://doi.org/10.1101/GR.215087.116

Kurtz, S., Phillippy, A., Delcher, A. L., Smoot, M., Shumway, M., Antonescu, C., & Salzberg, S. L. (2004). Versatile and open software for comparing large genomes. Genome Biology, 5(2), 1–9. https://doi.org/10.1186/GB-2004-5-2-R12/

Letunic, I., & Bork, P. (2019). Interactive Tree Of Life (iTOL) v4: recent updates and new developments. Web Server Issue Published Online, 47. https://doi.org/10.1093/nar/gkz239

Li, B., Cantino, P. D., Olmstead, R. G., Bramley, G. L. C., Xiang, C. L., Ma, Z. H., Tan, Y. H., & Zhang, D. X. (2016). A large-scale chloroplast phylogeny of the Lamiaceae sheds new light on its subfamilial classification. Scientific Reports 2016 6:1, 6(1), 1–18. https://doi.org/10.1038/srep34343

Li, H., & Durbin, R. (2009). Fast and accurate short read alignment with Burrows–Wheeler transform. Bioinformatics, 25(14), 1754–1760. https://doi.org/10.1093/BIOINFORMATICS/BTP324

Mader, M., Pakull, B., Blanc-Jolivet, C., Paulini-Drewes, M., Bouda, Z. H. N., Degen, B., Small, I., & Kersten, B. (2018). Complete chloroplast genome sequences of four meliaceae species and comparative analyses. International Journal of Molecular Sciences, 19(3). https://doi.org/10.3390/ijms19030701

Mark, J., & Rivers, M. C. (2017). Cedrela odorata, Spanish Cedar. The IUCN Red List of Threatened Species 2017, e.T32292A6, 10. http://dx.doi.org/10.2305/IUCN.UK.2017-3.RLTS.T32292A68080590.en

Michael, T. P., Jupe, F., Bemm, F., Motley, S. T., Sandoval, J. P., Lanz, C., Loudet, O., Weigel, D., & Ecker, J. R. (2018). High contiguity Arabidopsis thaliana genome assembly with a single nanopore flow cell. Nature Communications, 9(541). https://doi.org/10.1038/s41467-018-03016-2

Molinares, C., Prada, E., & León, E. de. (2019). CONDENANDO EL BOSQUE: Ilegalidad y falta de governanza en la Amazonia Colombiana.

Montero, M. I., Barrera, J. A., Benavides, B., & Lucena, A. (2016). Fichas técnicas de especies de uso forestal y agroforestal en la Amazonia Colombiana. Insituto Amazónico de Investigaciones Científicas SINCHI.

Paredes-Villanueva, K., de Groot, G. A., Laros, I., Bovenschen, J., Bongers, F., & Zuidema, P. A. (2019). Genetic differences among Cedrela odorata sites in Bolivia provide limited potential for fine-scale timber tracing. Tree Genetics and Genomes, 15(3). https://doi.org/10.1007/s11295-019-1339-4

Reiling, S. J., Chen, S.-H., & Ragoussis, I. (2020). McGill Nanopore Ligation LibPrep Protocol SQK-LSK109. Protocols. https://doi.org/dx.doi.org/10.17504/protocols.io.bpegmjbw

Schalamun, M., & Schwessinger, B. (2017). High molecular weight gDNA extraction after Mayjonade et al. optimised for eucalyptus for nanopore sequencing. Protocols. https://doi.org/dx.doi.org/10.17504/protocols.io.khkct4w

Schroeder, H., Cronn, R., Yanbaev, Y., Jennings, T., Mader, M., Degen, B., & Kersten, B. (2016). Development of Molecular Markers for Determining Continental Origin of Wood from White Oaks (Quercus L. sect. Quercus). PLOS ONE, 11(6), e0158221. https://doi.org/10.1371/JOURNAL.PONE.0158221

Scott, A. D., Zimin, A. V, Puiu, D., Workman, R., Britton, M., Zaman, S., Caballero, M., Read, A. C., Bogdanove, A. J., Burns, E., Wegrzyn, J., Timp, W., Salzberg, S. L., & Neale, D. B. (2020). A Reference Genome Sequence for Giant Sequoia. https://doi.org/10.1534/g3.120.401612

Shaw, J., Shafer, H. L., Rayne Leonard, O., Kovach, M. J., Schorr, M., & Morris, A. B. (2014). Chloroplast DNA sequence utility for the lowest phylogenetic and phylogeographic inferences in angiosperms: The tortoise and the hare IV. American Journal of Botany, 101(11), 1987–2004. https://doi.org/10.3732/AJB.1400398

Van der Auwera, G., & O’Connor, B. (2020). Genomics in the Cloud: : Using Docker, GATK, and WDL in Terra (O’Reilly Media (ed.); 1st Editio).

Wang, J., Liu, W., Zhu, D., Hong, P., Zhang, S., Xiao, S., Tan, Y., Chen, X., Xu, L., Zong, X., Zhang, L., Wei, H., Yuan, X., & Liu, Q. (2020). Chromosome-scale genome assembly of sweet cherry (Prunus avium L.) cv. Tieton obtained using long-read and Hi-C sequencing. Horticulture Research, 7(1). https://doi.org/10.1038/s41438-020-00343-8

Wang, W., Schalamun, M., Morales-Suarez, A., Kainer, D., Schwessinger, B., & Lanfear, R. (2018). Assembly of chloroplast genomes with long- and short-read data: A comparison of approaches using Eucalyptus pauciflora as a test case. BMC Genomics, 19(1), 1–15. https://doi.org/10.1186/S12864-018-5348-8

Wang, Y., Zhao, Y., Bollas, A., Wang, Y., & Au, K. F. (2021). Nanopore sequencing technology, bioinformatics and applications. Nature Biotechnology 2021 39:11, 39(11), 1348–1365. https://doi.org/10.1038/s41587-021-01108-x

Wei, S. J., Lu, Y. Bin, Ye, Q. Q., & Tang, S. Q. (2017). Population genetic structure and phylogeography of Camellia flavida (Theaceae) based on chloroplast and nuclear DNA sequences. Frontiers in Plant Science, 8, 718. https://doi.org/10.3389/FPLS.2017.00718/BIBTEX

Wöhner, T. W., Emeriewen, O. F., Wittenberg, A. H. J., Schneiders, H., Vrijenhoek, I., Halász, J., Hrotkó, K., Hoff, K. J., Gabriel, L., Lempe, J., Keilwagen, J., Berner, T., Schuster, M., Peil, A., Wünsche, J., Kropop, S., & Flachowsky, H. (2021). The draft chromosome-level genome assembly of tetraploid ground cherry (Prunus fruticosa Pall.) from long reads. Genomics, 113(6), 4173–4183. https://doi.org/10.1016/J.YGENO.2021.11.002

