## Supplementary figures and images for "New variants in the Chloroplast Genome Sequence of two Colombian individuals of the Cedar timber species (*Cedrela odorata* L.), using long read Oxford Nanopore Technology"

### Supplementary Figure S1

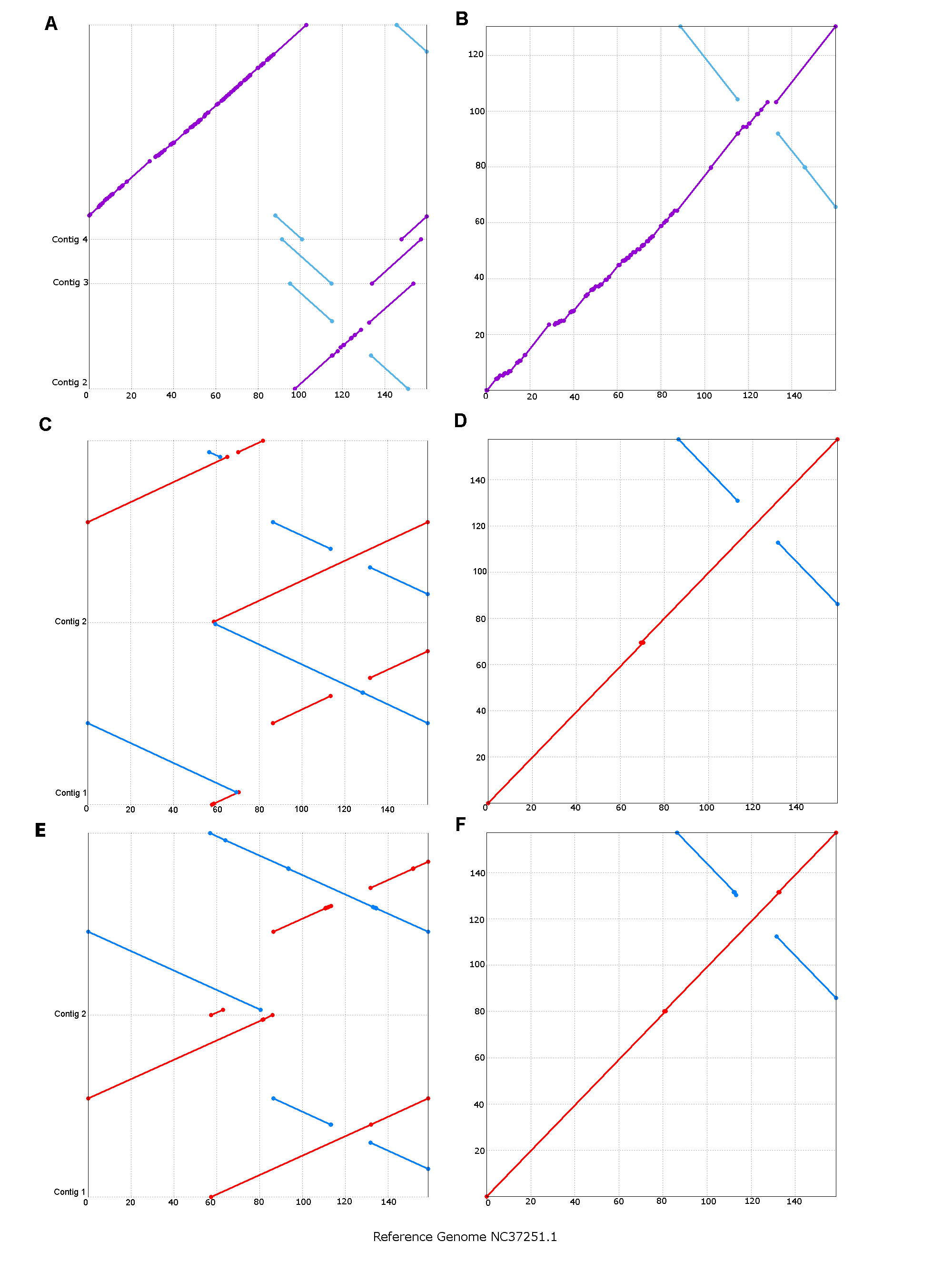
